# Characterizing the epigenetic signatures of the human regulatory elements: A pilot study

**DOI:** 10.1101/059394

**Authors:** Sawyer L Clement, Hani Z Girgis

**Author notes:** Email: Sawyer L. Clement -; Hani Z. Girgis.

## Abstract

**Background:** Chromatin modifications have provided promising clues on how cells that share the same copy of the genome can perform distinct functions. It is believed that enhancers and promoters are marked by a single chromatin mark each, H3K4me1 and H3K4me3, respectively. However, other studies have indicated that enhancers and promoters share multiple chromatin marks, including H3K4me1/2/3 and H3K27ac. Therefore, we asked whether the epigenetic signatures of these regulatory elements consist of a single mark or multiple marks. Repetitive regions, repeats, are usually ignored. However, we found, in public data, that repeats include about 25% of active enhancers. Thus, we asked how the epigenetic signatures of repetitive and non-repetitive enhancers differ. We studied the four marks in IRM90 (human lung fibroblast) and H1 (human embryonic stem cell).

**Results:** Our results show that enhancers and promoters are enriched significantly with the four marks, which form pyramidal signatures. However, the relative lengths of the marks are different. The promoter signature is directional; H3K4me2/3 and H3K27ac tend to be present downstream of the transcription start site; H3K4me1 tends to be present upstream. H1-specific enhancers have a similar signature to IRM90-specific enhancers; however, it is not the case for active promoters of the two cell types. Interestingly, inactive enhancers show a residual signature that resembles the signature of active enhancers. Finally, the epigenetic signature of enhancers found in repeats is identical to that of enhancers found in non-repetitive regions.

**Conclusions:** In this study, we characterized the epigenetic signatures of active and inactive enhancers (pyramidal) as well as active promoters (directional-pyramidal) in two cell types. These signatures consist of four chromatin marks that have been reported to be associated with enhancers and promoters. Interestingly, about one quarter of active enhancers are found in repeats. Active enhancers within repeats and those outside repeats have the same epigenetic signature. These results have great potential to change the way Molecular Biologists think of repeats, and to expand our understanding of gene regulation.

## Background

Epigenetic modifications, including DNA methylation and histone modifications, are the primary mechanism for long term regulation of gene expression. As all cells in an organism have the same genome, it falls to differential epigenetic landscapes to determine whether a given cell becomes a neuron, a melanocyte, or a T-cell [1]. While DNA methylation adds a methyl group directly to the DNA strand, histone modifications act on histones, chromatin proteins around which DNA is wrapped [2].

These histone/chromatin modifications include methylation, acetylation, ubiquitylation, phosphorylation, and others. Furthermore, these marks have different effects when attached to different histones and at different histone locations; H3K27ac (acetylation of histone 3 at lysine 27) is distinct from H3K18ac (acetylation of histone 3 at lysine 18). Histone modifications alter gene expression by disrupting chromatin organization and by recruiting or blocking the binding of non-histone proteins to DNA. These proteins may include transcription factors that regulate gene expression or proteins that further modify the chromatin [2].

While DNA hyper- and hypomethylation is commonly associated with gene under- and over-expression, the effects of specific chromatin modifications are less clearly understood. However, the scientific community has been making steady progress toward understanding the effects of chromatin marks.

Heintzman, et al. [3] reported the association of enhancers and promoters with single chromatin marks: H3K4me1 and H3K4me3, respectively. The pilot phase of the ENCODE project suggested that enhancers and transcription start sites (TSSs) have “inverse” chromatin patterns. Specifically, enhancers are reported to have high H3K4me1 and low H3K4me3, whereas TSSs are reported to have low H3K4me1 and high H3K4me3. Additionally, five marks, including H3K4me1/2/3, can be used for predicting the TSSs [4]. A third study concluded that H3K4me2 and H3K4me3 are characteristic marks of TSSs. In addition, the same study indicated that “H3K4me1 signal was low but showed some evidence of enrichment further downstream from the TSS than H3K4me2 and H3K4me3” [5]. In sum, the studies published in 2007 were not in agreement. Some studies suggested that enhancers are marked by the abundance of H3K4me1, whereas promoters are marked by the abundance of H3K4me3 (the single mark hypothesis). However, other studies suggested that H3K4me1/2/3 are present around promoter regions and H3K4me1/3 are present around enhancers (the multiple marks hypothesis).

A study by Pekowska, et al. [6] reported the association of H3K4me1/2/3 and the enhancers specific to the T-cell. Further, the ENCODE project reported that H3K4me1/2 are associated with enhancers and H3K4me1/2/3 are associated with promoters; in addition, H3K27ac marks active promoters and active enhancers [7]. At the end of 2012, it was reported that active enhancers and active promoters share H3K4me1/2/3 and H3K27ac.

In 2013, a study by Zhu, et al. [8] suggested that H3K27ac and H3K4me3, among other chromatin marks, are indicative of active enhancers. Moreover, another study confirmed that H3K4me1/2/3 and H3K27ac are the best four chromatin marks indicative of active enhancers [9].

The multiple marks hypothesis is supported by a larger number of studies than the single mark hypothesis. Nonetheless, the single mark hypothesis is still cited in recent studies [10, 9].

Motivated to resolve these discrepancies, we conducted a study on publicly available data. The main purpose of our study is to characterize the epigenetic signatures of enhancers and promoters. Specifically, we studied the distributions of H3K4me1/2/3 and H3K27ac relative to the site and each other with three main questions in mind. First, which of the two hypotheses is more accurate? Second, how does the epigenetic signature of enhancers differ from the signature of promoters? Third, how does the epigenetic signature of enhancers found in repetitive regions (about 25% of active enhancers in the studied cells) differ from that of enhancers outside these regions?

## Results and Discussion

This research was conducted to characterize the epigenetic signatures of the main regulatory elements. We focused on the following four chromatin marks: H3K4me1/2/3 and H3K27ac. In this study, we examined the distribution of these marks in the context of promoters and enhancers, with the intention of determining a characteristic epigenetic signature for each.

### IMR90-specific enhancers exhibit pyramidal epigenetic signature

We began our investigation by examining the epigenetic signatures around active enhancers (p300 binding sites overlapping DNase I hypersensitive sites (DHSs) provided by another study by Rajagopal, et al. [9]). We chose the IMR90 cell line (human lung fibroblast) for our experiments, as the relevant epigenetic information for this tissue type is publicly available. We began by plotting the distribution of the chosen marks about individual active enhancers (Figures 1a-1f). Eventually, we detected a pattern in the epigenetic distribution, more complex than the simple presence or absence of particular marks. Stretches of each of the four marks appeared in most of the plots, approximately centered about the DHSs. The overlapping stretch of H3K4me1 was usually the longest, followed by H3K4me2 and H3K27ac (roughly equal in length), with the H3K4me3 usually being the shortest. When plotted about a single enhancer, we observed that the four marks form a pyramidal shape. These results confirm that the enhancer epigenetic signature is not defined by the presence or absence of H3K4me1 (the single mark hypothesis), but by the arrangement of the four marks around the enhancer (the multiple marks hypothesis).

**Figure 1:**
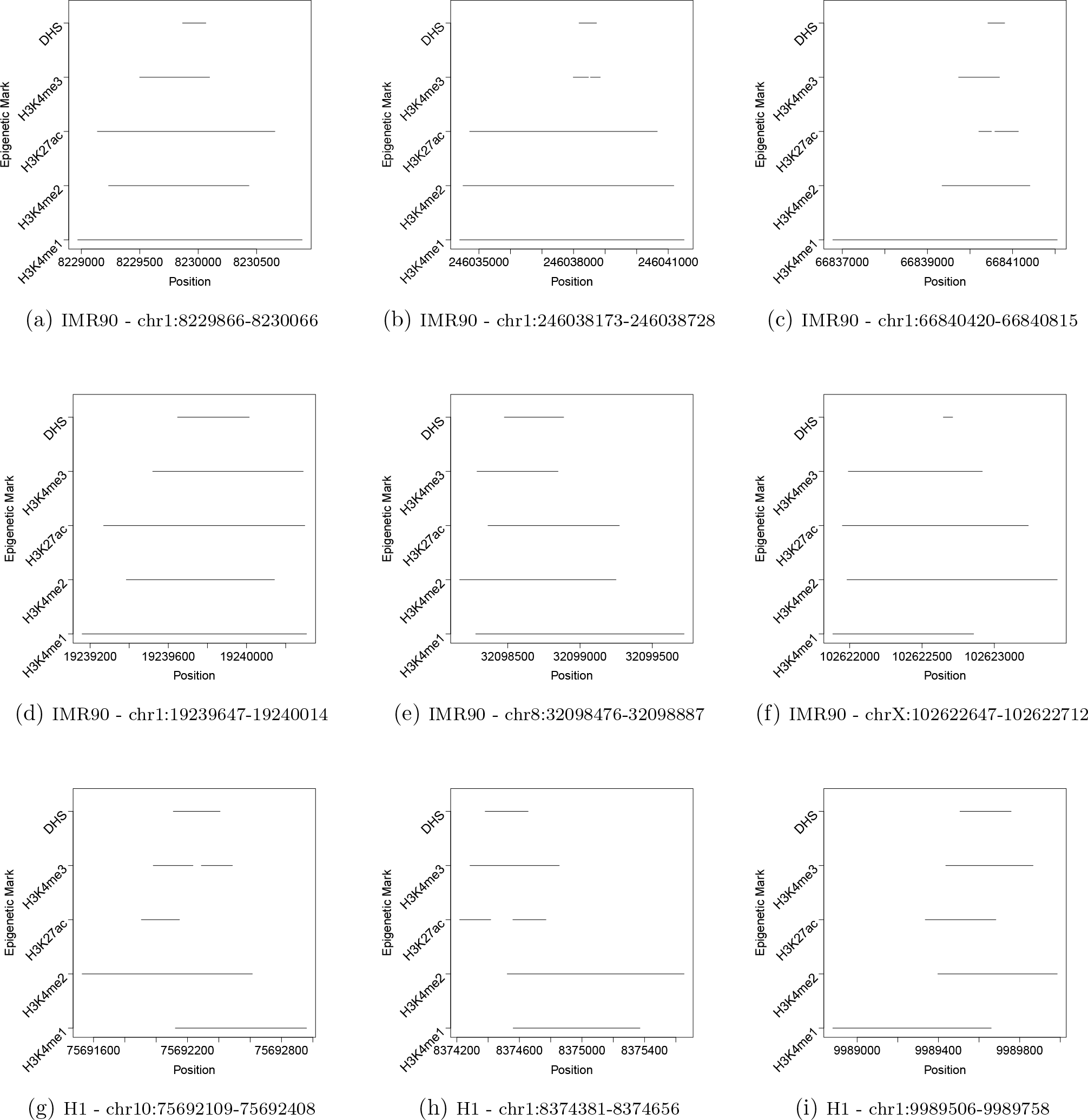
The arrangement of H3K4me1/2/3 and H3K27ac around active enhancers specific to the IMR90 and the H1 cell lines. Active enhancers are defined as p300 binding sites overlapping DNase I hypersensitive sites (DHSs). All coordinates are according to the hg19 assembly. (a-f) The four chromatin marks form pyramidal shape about enhancers specific to IMR90. (g-i) The four marks form stacked/pyramidal shape about enhancers specific to H1. The arrangement of the four marks around the IMR90-specific enhancers is similar, though not identical, to that around the H1-specific enhancers.

Next, we sought to determine whether the pyramidal enhancer pattern would persist in a large data set. However, we had no reference for how these signatures compare to the epigenome as a whole. Therefore, we examined the distributions of the four marks around 500 segments (each is 500 bp long), spread uniformly throughout the human chromosome 1. We refer to these sequences as control sequences. These random segments have low content of the epigenetic marks, though the actual content varied somewhat between the mark types. H3K4me1 and H3K4me2 appeared in approximately 30% of samples, whereas H3K27ac and H3K4me3 appeared in approximately 15%.

The epigenetic signatures of 2000 active enhancers were profiled (Figure 2). We found that each of the four chromatin marks was consistently present at active enhancers (80-95%). Further, the enhancer regions were 3.1-fold more enriched with H3K4me1 than the control sequences (P-value < 2.2e^−16^, Fisher’s exact test). Similarly, the other three marks were significantly enriched in the enhancer regions (H3K4me2: 3.4 folds, H3K4me3: 5.2 folds, H3K27ac: 5.2 folds; P-value < 2.2e^−16^, Fisher’s exact test). Again, these results show that the four marks are enriched in the enhancer regions.

**Figure 2:**
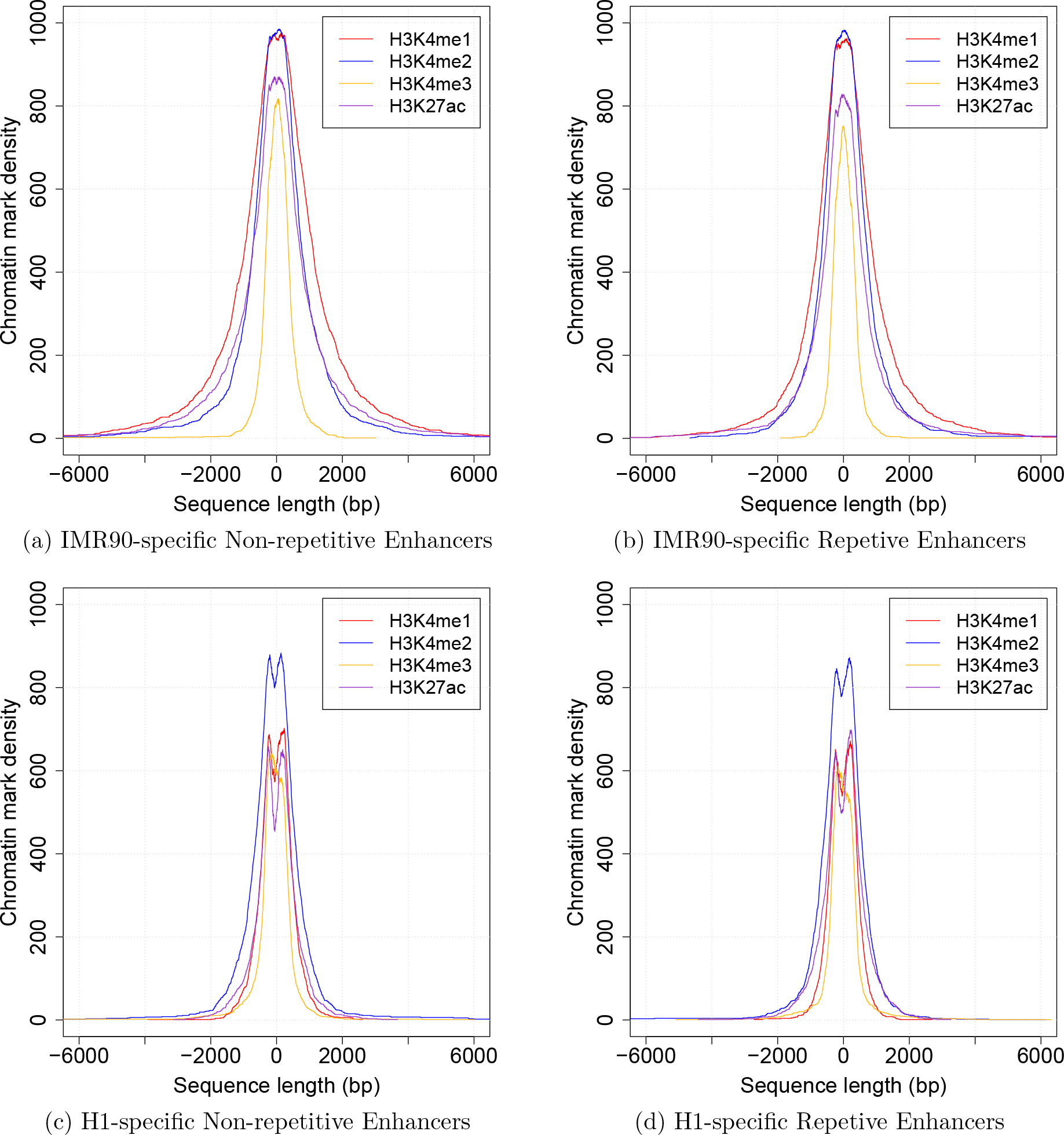
Repetitive (overlapping repetitive elements) and non-repetitive (outside repetitive elements) enhancers exhibit identical epigenetic signatures. Graphs show summed chromatin mark densities of 1000 enhancers centered around the p300 binding sites. (a & b) Comparisons of chromatin mark densities within the repetitive and the non-repetitive IMR90-specific enhancers. The repetitive and the non-repetitive IMR90-specific enhancers conform strongly to the pyramidal epigenetic signature. (c & d) Comparisons of chromatin mark densities within the repetitive and the non-repetitive H1-specific enhancers. The repetitive and the non-repetitive H1-specific enhancers have identical stacked epigenetic signatures.

### Epigenetic Marks of Enhancers in repetitive and non-repetitive regions

Interestingly, H3K4me3 was more enriched in the enhancers than was H3K4me1 (5.2 folds vs. 3.1 folds). This observation questions the common assumption that H3K4me1 is the main chromatin mark characterizing enhancers. This assumption is based on the abundance of the mark, not on the enrichment value obtained by comparing the observed abundance in the enhancers to that in the control sequences.

The width and the orientation patterns observed in the individual enhancer figures also reappeared. H3K4me1 had the largest average width (5000 bp), followed by H3K4me2 and H3K27ac (3000-4000 bp), with H3K4me3 being the narrowest (1500-2000 bp). Together, these observations support the pyramidal distribution of the studied epigenetic marks about the active IMR90-specific enhancers.

### Active enhancers in repetitive regions exhibit the same epigenetic pattern as active enhancers outside repetitive regions

A study by Xie et al. [10] conducted on a number of tissues showed (i) transposon subfamilies have different patterns of hypomethylation across tissue types; (ii) these “differentially-hypomethylated” subfamilies are associated with H3K4me1; (iii) they are associated with the expression of genes in their vicinities; and (iv) the sequences of these subfamilies include binding sites for tissue-specific transcription factors. These four findings suggest that transposon subfamilies have tissue-specific enhancer-like functions. Motivated by these findings, we divided active IMR90 enhancers into those overlapping with repetitive regions and those that are not. Out of 25,109 enhancers, 5,925 (23.6%) overlap repetitive regions. We wished to determine whether, if the signature persisted, it would appear on both repetitive and non-repetitive enhancers. Therefore, we profiled the chromatin marks around 1000 non-repetitive enhancers and 1000 repetitive enhancers (Figures 2a and 2b). These figures show that there are no differences between the distribution of these four marks on repetitive and non-repetitive enhancers, indicating that non-repetitive and repetitive enhancers have the same epigenetic signature.

### The four epigenetic marks are significantly depleted in inactive enhancers compared to active enhancers

We had confirmed that the epigenetic signature of enhancers can be observed in both individual enhancers and large enhancer sets. However, we had not performed any large-scale analysis of inactive enhancers. As such, we could not be certain whether the pyramidal signature observed previously was indicative only of active enhancers or of enhancers as a whole. To resolve this, we compared the epigenetic signatures of an equal number of inactive and highly active enhancers, as determined by eRNA (enhancer RNA) levels (obtained from the Fantom5 project [11]). Figures 3a and 3b show the epigenetic signatures of the active and the inactive enhancers. Active enhancers were significantly more enriched with the four marks than inactive ones (H3K4me1: 1.5 folds, P-value = 0.000259; H2K4me2: 1.6 folds, P-value = 3.035e^−7^; H3K4me3: 2.8 folds, P-value = 1.677e^−15^; H3K27ac: 2.0 folds, P-value = 1.064e^−8^; Fisher’s exact test). Additionally, the marks were clearly narrower in the inactive enhancers than those found in the active enhancers. Overall, the inactive enhancers displayed significantly decreased levels of all four epigenetic marks, indicating that the enhancer epigenetic signature is weaker in these enhancers.

**Figure 3:**
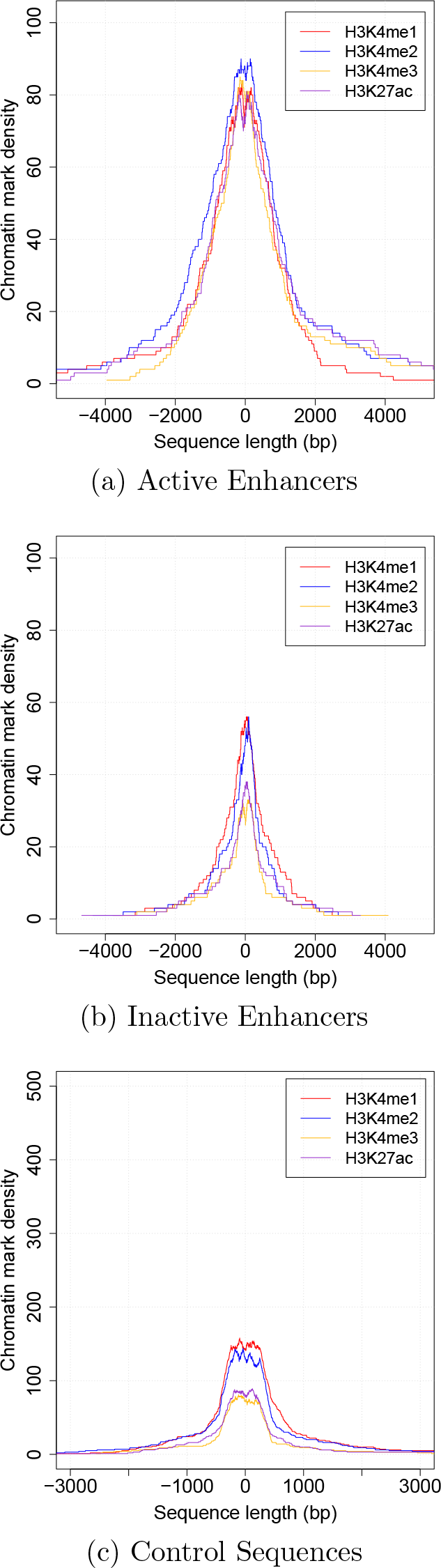
Comparisons of chromatin mark densities within (a) active enhancers specific to IMR90, (b) inactive enhancers that are active in other cells, and (c) the control sequences. Chromatin marks around the active enhancers are both broader and more common than those around the inactive enhancers. The inactive enhancers are significantly more enriched with the four marks than the control sequences.

### Inactive enhancers show residual enrichment of the four epigenetic marks, resembling the pyramidal signature

Next, we asked whether the densities of the marks in inactive enhancers are similar to the genome average, i.e. similar to the control sequences. Therefore, we compared these densities in the inactive enhancers and the control sequences (Figures 3b and 3c). Interestingly, inactive enhancers were more enriched with the four marks than the control sequences (H3K4me1: 1.8 folds, P-value = 9.838e^−6^; H2K4me2: 1.9 folds, P-value = 1.289e^−6^; H3K4me3: 2.0 folds, P-value = 0.0007712; H3K27ac: 2.5 folds, P-value = 3.752e^−7^; fisher’s exact test). These results suggest that inactive enhancers exhibit a weaker, yet significant, version of the epigenetic signature of the tissue-specific active enhancers.

### The H1-specific enhancers and the IMR90-specific enhancers have similar epigenetic signatures

The study by Rajagopal et al. [9] determined enhancers specific to the H1 cell line experimentally (p300 binding sites overlapping DHSs). We asked if the epigenetic signature observed in the enhancers specific to the IMR90 is the same/or similar to that of the enhancers specific to H1. Figures 1g-1i show the four marks around three H1-specific enhancers. The four marks are stacked around the enhancers, suggesting a stacked/pyramidal epigenetic signature.

Next, we profiled the epigenetic signature of 2000 H1-specific enhancers (Figure 2). The four studied marks are present around the active enhancers of H1. These profiles differ from those of IMR90 in the width of the densities and in the relative order of the bottom two layers. In general, the densities observed in H1 are narrower that those observed in IMR90. Recall that the signature of the IMR90-specific enhancers consists of these layers: H3K4me1 (the widest), H3K4me2 and H3K27ac (roughly the same width), and H3K4me3 (the narrowest). The signature of the H1-specific enhancers consists of these layers: H3K4me2 (the widest), H3K4me1 and H3K27ac (roughly the same width), and H3K4me3 (the narrowest). The two signatures differ in the relative order of the lower two layers (H3K4me1 and H3K4me2). These results show that the epigenetic signatures of the H1-specific enhancers and the IMR90-specific enhancers are similar, though not identical.

### H1-specific enhancers in repetitive regions exhibit the same epigenetic signature as those outside repetitive regions

We observed that 23.8% (1,402 out of 5,899) of the H1-specific enhancers overlap repetitive regions. Recall that a similar percentage of the IMR90-specific enhancers overlap repetitive regions as well. Furthermore, the epigenetic profiles of 1000 non-repetitive enhancers and 1000 repetitive enhancers of the H1 are almost identical (Figures 2c and 2d). These results confirm the results observed in the non-repetitive enhancers and the repetitive enhancers of IMR90.

Given the existence of a pyramidal pattern about active enhancers, our next consideration was whether other regulatory regions, namely promoters, might exhibit a similar signature.

### Active promoters exhibit directional-pyramidal epigenetic signature

We examined the epigenetic signature of individual active promoters. First, we defined active promoters as transcription start sites (TSSs) overlapping DHSs. However, a few problems prevented us from receiving immediate results. Initially, we drew random DHS-overlapping promoters from a list of all known TSSs. However, as gene expression in a cell varies greatly over time, not necessarily coinciding with DHS establishment, the resulting figures were inconsistent. Despite this, a few promoters did appear to demonstrate a directional-pyramidal pattern. Therefore, we decided to repeat the experiment with a more precise method for determining active promoters based on gene expression levels.

The second time, we chose ten promoters of genes with the highest expression (active promoters) in IMR90 as well as ten promoters of unexpressed genes (inactive promoters). The epigenetic plots made for the inactive promoters showed no apparent pattern, while each plot made for the active promoters demonstrated some version of the directional-pyramidal pattern observed in the first trial (Figures 4a-4c). As with the pattern observed about enhancers, the pattern about promoters involved all four of the studied chromatin marks (H3K4me1/2/3 and H3K27ac). The distinctions were in (i) the order of the layers of the signature and (ii) how these marks were arranged around the TSS, i.e. the directionality.

**Figure 4:**
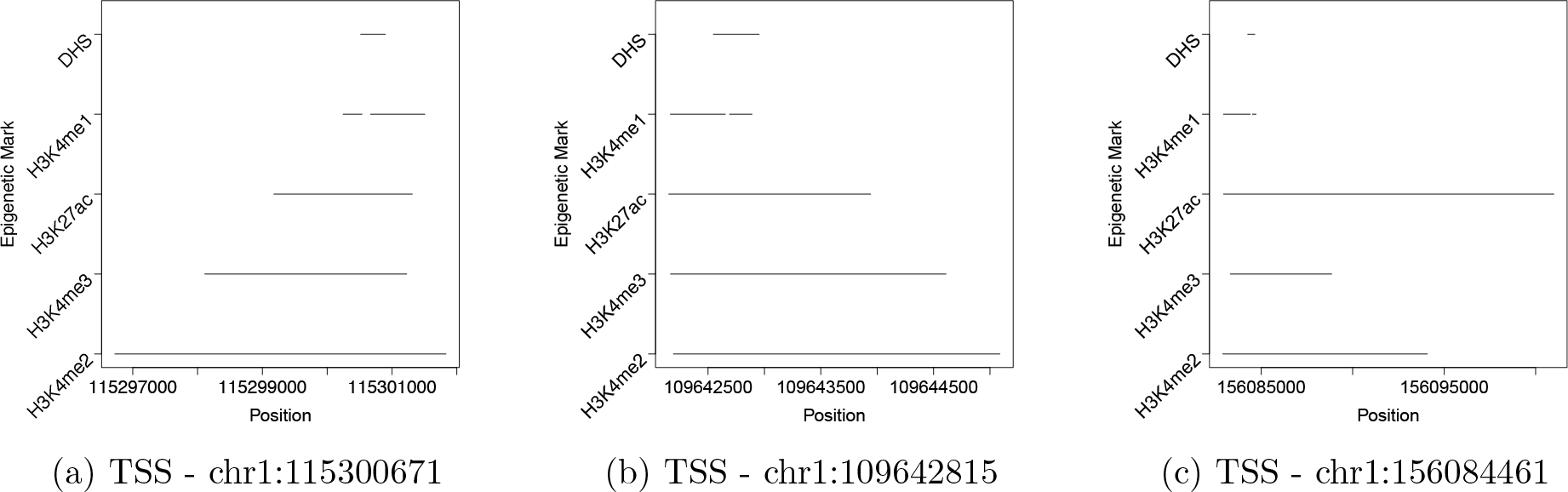
The epigenetic signature of active promoters in IMR90. (a-c) Four chromatin marks (H3K4me1/2/3 and H3K27ac) form directional pyramidal shape about three promoters of highly expressed genes in the IMR90 cell line. All coordinates are according to the hg19 assembly.

The layers of the directional-pyramidal signature of the active promoters were: H3K4me2 (the broadest), H3K4me3, H3K27ac, and H3K4me1 (the narrowest). Recall that the layers of the pyramidal signature of the active enhancers were: H3K4me1 (the broadest), H3K4me2, H3K27ac, and H3K4me3 (the narrowest).

Additionally, we found that H3K4me2/3, and H3K27ac all encompassed the TSS-overlapping DHS. Moreover, they were each more present downstream of the promoter than upstream. In contract, H3K4me1 was mainly present upstream of the DHS, with a small break in the region overlapping the DHS (possibly at the site of the TSS itself). The orientation of the chromatin marks relative to the DHS was inverted on positive strand promoters compared to negative strand promoters.

Having observed a distinctive epigenetic pattern in individual promoters, our next step was to determine whether this pattern would appear in a large data set consistently. To this end, we studied the promoters (+/− 250 bp from TSS) of the 100 most expressed (active promoters) and 100 unexpressed genes (inactive promoters) as determined by RNA-seq values.

We compared the four epigenetic marks across the two sets. The active promoters showed clear enrichment of these marks compared to the inactive promoters (H3K4me1: 1.9 folds, P-value = 8.826e^−07^; H3K4me2: 2.3 folds, P-value = 5.033e^−14^; H3K4me3: 3.6 folds, P-value < 2.2e^−16^; H3K27ac: 3.6 folds, P-value < 2.2^e^ −16; Fisher’s exact test).

As expected, active promoters were also enriched with the four marks compared to the control sequences (H3K4me1: 2.4 folds, P-value = 3.903e^−16^; H3K4me2: 3.1 folds, P-value < 2.2e^−16^; H3K4me3: 6.0 folds, P-value < 2.2e^−16^; H3K27ac: 5.6 folds, P-value < 2.2e^−16^; Fisher’s exact test).

H3K4me1 around active promoters demonstrated a clear drop at the TSS (Figure 5a). This drop mirrored the H3K4me1 TSS breaks observed in the individual promoters.

**Figure 5:**
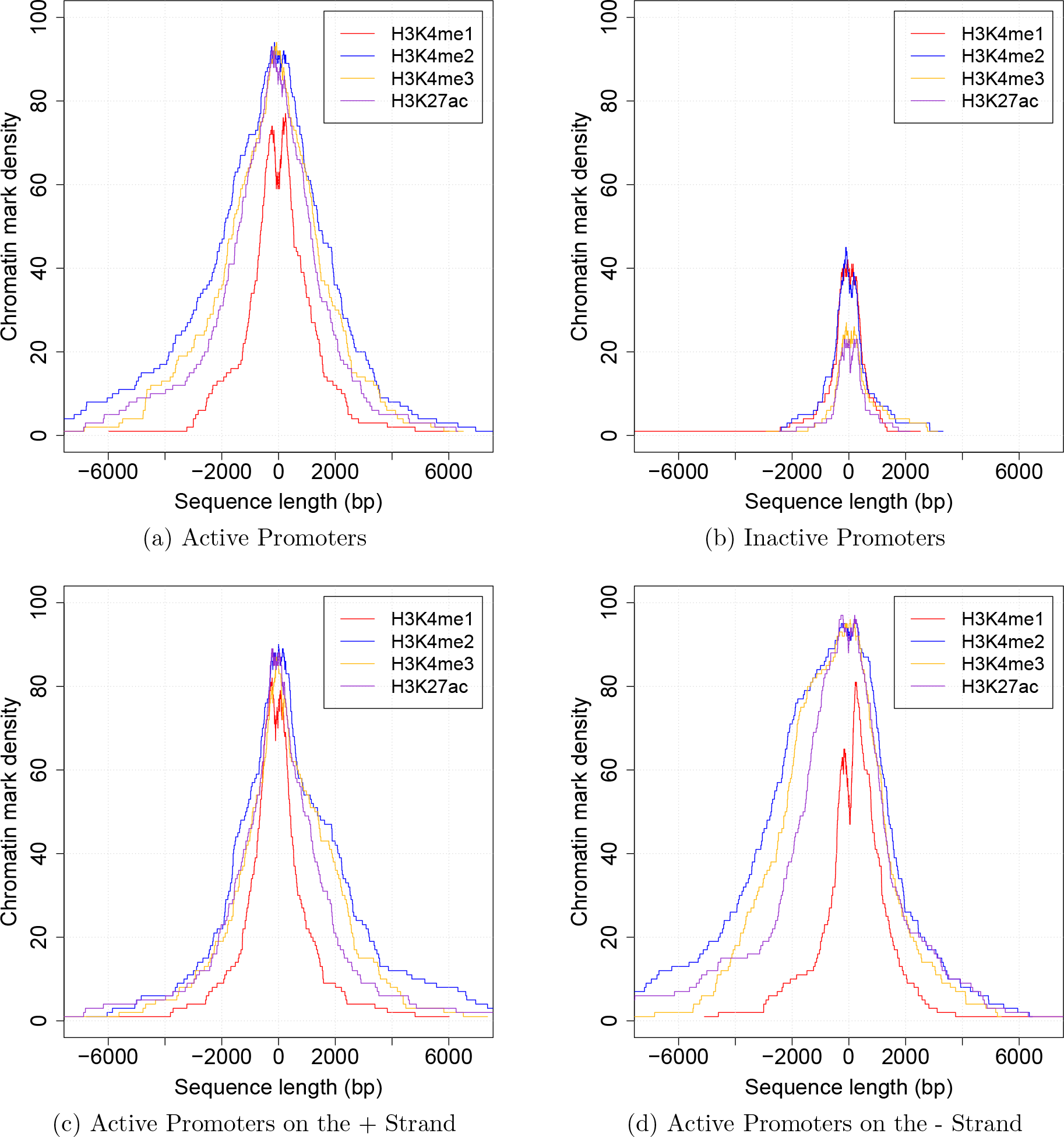
Active promoters in IMR90 exhibit directional-pyramidal epigenetic signature. Each graph shows summed chromatin mark density of 100 promoters centered around the TSS. (a & b) Comparisons of chromatin mark densities between the active (promoters of the 100 most expressed genes) and the inactive promoters (promoters of unexpressed genes in IMR90). Active promoters are significantly enriched with all marks studied. (c & d) Comparisons of chromatin mark densities within the active promoters on the negative and the positive strands. Epigenetic patterns of opposite strands roughly mirror each other, corresponding to opposite directions of transcription. H3K4me2, H3K4me3, and H3K27ac are enriched downstream of the promoter, while H3K4me1 is enriched upstream.

The individual promoter regions show that certain epigenetic marks extend farther on one side of the TSS than the other. The directions of these marks suggest that their distributions are related to the direction of transcription. Therefore, we analyzed the active promoters on the positive and the negative strands separately. Figures 5c and 5d show the epigenetic marks of the 100 active and the 100 inactive promoters. The results matched the epigenetic signature observed in the individual promoters. The directionality was reversed across opposing strands (positive and negative), indicating that the pattern is related to transcription direction. On both strands, H3K4me2/3 and H3K427ac tended to be enriched downstream of the promoter. H3K4me1 tended to be enriched upstream, with the gap at the TSS appearing once again.

### Unlike inactive enhancers, the marks around the inactive promoters does not resemble the directional-pyramidal signature

Inactive promoters are weakly enriched with three of the four marks (H3K4me2: 1.4 folds, P-value = 0.03304; H3K4me3: 1.7 folds, P-value = 0.01846; H3K27ac: 1.6 folds, P-value = 0. 04251; Fisher’s exact test). However, the densities of these marks do not resemble those of the active promoters, supporting the notion that promoter epigenetic signature is related to gene activation.

### Active promoters of the H1 cell line are not marked epigenetically

We studied the active promoters of the H1 cell line using the same procedure used in studying the IMR90 active promoters. However, the directional-pyramidal pattern was not observed in the H1 cell line. Moreover, the densities of the marks around active promoters resemble the genome average. These observations may be due to H1 being a stem cell, as they are known to have a “unique epigenetic signature” [12].

### Discussion

In this study, we characterized the epigenetic signatures of enhancers and promoters in the IMR90 and the H1 cell lines. We were motivated by the discrepancies in the literature about how certain chromatin marks are associated with these regulatory elements. Some studies report that H3K4me1 marks enhancers, whereas H3K4me3 marks promoters. Yet other studies report that four chromatin marks (H3K4me1/2/3 and H3K27ac) are present around enhancers and promoters. The first question we considered in this study whether the epigenetic signatures of promoters and enhancers consist of one mark or multiple marks. Our analyses show that these signatures consist of multiple marks, including, but not limited to, all of the four marks studied. We then asked how the epigenetic signature of the promoters differs from the enhancer signature. Additionally, we asked how the signature of active enhancers found in repetitive regions differs from the signature of the ones outside repetitive regions. In sum, this study contributes the following seven findings:

- In the IMR90 cell line, H3K4me1/2/3 and H3K27ac form a pyramidal shape around active enhancers. H3K4me1 is the base of the pyramid. H3K4me2 and H3K27ac are the middle layers. H3K4me3 is the top of the pyramid. The regularity with which this pattern appears suggests that it is intrinsically tied to transcription in this cell line.
- Active enhancers specific to IMR90 are more enriched with H3K4me3 than with H3K4me1 (5.2 folds vs. 3.1 folds), questioning the common assumption that H3K4me1 is the main chromatin mark characterizing active enhancers. This assumption is supported by the abundance, not the enrichment value relative to the genome average, of H3K4me1 around enhancer regions.
- Active promoters of IMR90 demonstrate a directional-pyramidal epigenetic signature. H3K4me2 is the base of the pyramid. H3K4me3 and H3K27ac are the middle layers. H3K4me1 is the top of the pyramid. Note that the directional-pyramidal signature of active promoters is roughly the “inverse” of pyramidal signature of active enhancers with regard to chromatin mark length.
- Chromatin marks in active IMR90 promoters are unevenly distributed about the TSS. H3K4me2/3 and H3K27ac tend to be more present downstream of the TSS, whereas H3K4me1 tends to be more present upstream, and often has a characteristic gap around the TSS. This transcription-dependent directionality around the TSS suggests that these marks are involved with the initiation of transcription.
- The epigenetic signature of the enhancers active in H1 (embryonic stem cell) is similar to the signature of those active in IMR90. However, the promoters of genes active in H1 do not show any recognizable epigenetic pattern consisting of the four studied marks.
- Inactive enhancers in IMR90 exhibit a residual epigenetic signature that resembles the signature of active enhancers, whereas inactive IMR90 promoters do not exhibit any such signature.
- The epigenetic signatures of active enhancers in non-repetitive regions and those in repetitive regions are indistinguishable. As this signature is linked to enhancer activation, this reinforces the notion that repetitive elements have significant regulatory function. As such, we urge the scientific community to stop masking/ignoring repeats and to start studying them.

## Materials and Methods

### Data

In this study, we used enhancers experimentally determined by Rajagopal et al. [9]. An enhancer is defined as a DNase I hypersensitive site (DHS) where p300 binds “distal to known UCSC and Gencode” transcription start sites (TSSs). Enhancers studied are specific to the IMR90 cell line (human lung fibroblasts) and the H1 cell line (human embryonic stem cells). Chromatin modification data sets, including those for H3K4me1/2/3 and H3K27ac, were downloaded from the publicly available Human Epigenome Atlas [13]. The DHS data sets were downloaded from the NCBI Gene Expression Omnibus, under the designation Sample GSM468792 [9]. RNA-seq data were downloaded from the ENCODE project [14]. Name, TSS, and chromosome location data for all human genes were downloaded from the Ensembl website [15]. As the Ensembl data are in hg38 and the other data sets are in hg19, the Ensembl data were converted into hg19 using the UCSC LiftOver tool. Repeats of the human genome (hg19) detected by RepeatMasker were downloaded from the Institute for Systems Biology website [16]. The eRNA (enhancer RNA) levels for the fetal lung tissue were downloaded from the FANTOM5 project [17, 18, 11].

### Individual Enhancers

Individual enhancers (p300 binding sites overlapped by DHSs) were manually selected. For each p300 binding site, the overlapping DHS was located. Then, segments representing the four chromatin marks that overlapped the DHS were detected. These chromatin mark segments were then plotted one on top of another as lines to easily determine how these marks were distributed about the DHS (and by extension the enhancer).

### Multi-Enhancers

Active enhancers were taken from a list of experimentally determined enhancers (p300 binding sites overlapping DHSs and distal to known TSSs) [9]. Using a list of repetitive elements in the human genome, these enhancers were separated into two sets: those found in repetitive elements and those found in non-repetitive regions. The first 1000 enhancers from each set were arbitrarily chosen. For each chromatin mark, the length and the distribution about an enhancer were recorded. Then segments representing these marks were summed and plotted together in order to display the distribution of that mark around the enhancers. Separate analyses were done with repetitive and non-repetitive enhancers for purposes of comparison.

### Control Sequences

We constructed a set of control sequences by selecting 500 segments distributed uniformly throughout the human chromosome one. Each segment is 500 bp long. The IMR90 chromatin marks overlapping the control sequences were analyzed and summed as done previously.

### Active and Inactive Enhancers

The eRNA (enhancer RNA) data was used for selecting a set of enhancers highly active in IMR90 and an equal-sized set of enhancers inactive in IMR90. All enhancers with eRNA expression values 19 or greater were chosen as active enhancers. This cutoff value was chosen because it yielded approximately 100 (102) of the most active enhancers, enough to clearly demonstrate any major epigenetic pattern. The set of inactive enhancers consisted of 102 enhancers with eRNA expression values of 0. The chromatin marks around each enhancer set were analyzed as was done previously. Because these enhancers were not single coordinates (as were the p300 binding sites) but rather representative regions, all chromatin marks overlapping these regions were recorded.

### Individual Promoters

Initially, individual promoters were manually selected from a list of all human promoters. These promoters were analyzed the same way as the individual enhancers, plotting the epigenetic marks around DHS-overlapping promoters. When this provided an unclear pattern, manually chosen promoters were replaced with ten promoters of the most expressed genes in a specific cell line. Additionally, ten promoters of unexpressed genes in the same cell line were selected. The chromatin marks overlapping the promoters were plotted and analyzed as done previously.

### Multi-Promoters

We determined the 100 most expressed genes and 100 unexpressed genes in a specific cell line based on the ENCODE gene expression data. As the gene expression data (ENCODE) and the TSS data (Ensembl) were from different sources, some gene names from one did not appear in the other; such genes were removed from consideration. Because the gene expression data were sufficient proof of promoter activation, the requirement for a promoter to overlap a DHS was removed. Instead, all chromatin marks near the promoters (+/− 250 base pair from the TSS) were included in the analysis. For each chromatin mark, the length and the distribution about the promoters of the studied genes were recorded. The chromatin marks around the promoters were analyzed as was done with the enhancers. Separate analyses were done for the 100 most expressed and the 100 unexpressed genes, to contrast the distribution of marks in inactive and active promoters. This procedure was repeated with the additional consideration of gene strand. This time, the 100 most expressed and unexpressed genes on the positive strand and the negative strand were analyzed separately.

## List of abbreviations

TSS: transcription start site; DHS: DNase I hypersensitive site; eRNA: enhancer RNA.

## Declarations

### Ethics approval and consent to participate

Not applicable.

### Consent for publication

Not applicable.

## Availability of data and material

All data sets used in this study are publicly available as noted under the Materials and Methods Section.

## Authors' contributions

SLC designed and implemented the analyses. HZG initiated the study. HZG designed the analyses and calculated the statistical tests. SLC and HZG wrote the manuscript. All authors read and approved the final manuscript.

## Acknowledgements

Not applicable.

## Authors’ information

SLC is a senior student at the University of Tulsa. His major is Biology. HZG is an Assistant Professor of Computer Science at the University of Tulsa. HZG did his postdoctoral work at the National Center for Biotechnology Information (NCBI), the National Institutes of Health (NIH). HZG has been studying regulatory elements and repeats for seven years. HZG majored in Biology and Computer Science while he was undergraduate student at the State University of New York at Buffalo.

